# Spatiotemporal Control of Genetic and Epigenetic Editing through Covalent Tethering of CRISPR Nanoparticles to Zwitterionic Microgels

**DOI:** 10.64898/2026.09.18.751510

**Authors:** Joshua P. Graham, Erika E. Wheeler, J. Kent Leach, Tomas Gonzalez-Fernandez

## Abstract

CRISPR gene editing offers unprecedented genomic and transcriptomic control, positioning it as a powerful tool for cell therapies. Non-viral CRISPR delivery avoids the immunogenicity, genomic integration, and packaging limits of viral vectors, but local or systemic injections of non-viral nanoparticles suffers from transient action and poor biodistribution. Alternatively, biomaterial-based delivery improves nanoparticle localization and sustains delivery, yet current approaches rely on non-specific adsorption of nanoparticles to scaffolds, risking aggregation, destabilization, and unreliable release kinetics. This work establishes a novel strategy to covalently tether nanoparticles to biomaterial substrates through SnoopTag and SpyTag bioconjugation systems for spatiotemporal control of non-viral CRISPR delivery. The RALA cell-penetrating peptide, an efficient and low-cytotoxicity CRISPR delivery system, was functionalized with SnoopTag without altering nanoparticle formation or transfection capacity. SnoopTag-functionalized nanoparticles were then covalently tethered to SpyTag-decorated zwitterionic microgels via a SnoopCatcher-SpyCatcher fusion protein. This platform achieves sustained RALA-mRNA nanoparticle delivery and prolonged CRISPR activation in human mesenchymal stem cells seeded within CRISPR-loaded microgel scaffolds. This work establishes a defined biorthogonal conjugation system that tethers non-viral nanoparticles to biomaterial platforms for sustained and localized CRISPR gene editing.

## 1. Introduction

CRISPR gene editing offers a precise and tunable platform for genetic and epigenetic regulation.^1^ These systems use a guide RNA (gRNA) to direct a Cas endonuclease to specific target genes, enabling precise gene editing outcomes including gene knock-in, knock-out, knock- down, transcriptional activation or inhibition, base editing, and prime editing to benefit a range of therapeutic applications.^2–12^ However, CRISPR delivery in therapeutic contexts remains a significant challenge. The first FDA-approved CRISPR therapy, Casgevy, uses electroporation for reliable CRISPR delivery and editing of hematopoietic stem cells (HSCs) *ex vivo*.^13^ However, electroporation is limited by trade-offs between editing efficiency and cell viability and cannot be used for *in vivo* or *in situ* gene editing, limiting its therapeutic scope and increasing the costs and complexity of biomanufacturing pipelines.^14^ Alternatively, viral vectors through adeno-associated viral (AAV) or lentiviral systems can reliably delivery CRISPR machinery for constitutive gene editing *in vivo*, but they are limited by immunogenicity and genomic integration risks associated with adverse clinical events such as tumorigenicity and death.^15–18^ Instead, non-viral lipid nanoparticle- (LNP)-based delivery is receiving increased attention due to its use for the COVID-19 vaccine.^19,20^ However, LNPs activate inflammatory pathways, serving as self-adjuvants to ensure immune activation.^21^ While this immune activation is beneficial for vaccines and viral immunity, it can hamper the therapeutic potential of CRISPR gene editing in applications such as regenerative medicine, where inflammation-driven catabolic pathways can disrupt regenerative procresses.^22^ To address these limitations, multiple non-viral alternatives have been developed to replace LNP-based gene delivery.^23^ We have previously optimized the arginine-alanine-leucine-alanine (RALA) cell-penetrating peptide (CPP) for efficient CRISPR gene editing in primary human mesenchymal stem cells (MSCs) with reduced immunogenicity compared to lipid- and cationic polymer-based non-viral systems.^14^ RALA-based CRISPR delivery also achieved efficient CRISPR knock-in, knock-out, and epigenetic regulation in MSCs via plasmid DNA, RNA, or ribonucleoprotein (RNP) delivery.^24^ Despite the advantages of RALA-based CRISPR delivery, its therapeutic application is limited by poor biodistribution after local injection, accumulation in the lungs and liver after systemic administration, and transient gene delivery, prompting the need for spatiotemporal control systems to increase on target editing efficiencies while avoiding editing in non-target organs.^25^

Non-viral nanoparticles are typically administered via systemic or local injections which fail to achieve effective editing rates in target tissues.^20,24,26^ Following systemic administration, nanoparticles rapidly accumulate in the liver, spleen, or lungs based on their physiochemical properties.^25,27–29^ Although specific cell-targeting nanoparticles have been designed to improve editing rates at specific locations, these primarily target liver, lung, or tumor cells, and are inadequate for the majority of biological systems.^30–35^ Local injections can improve nanoparticle delivery to target tissues, but nanoparticles are still rapidly cleared from the injection site, limiting editing rates and risking off-target editing. In fact, multiple studies have reported low gene editing efficiencies less than 15% in the muscle, brain, liver, and lungs following hydrodynamic injections due to poor nanoparticle distribution and limited duration of gene editing.^36,38,40^ To overcome these low editing rates and achieve reliable therapeutic effects, repeated injections are required, leading to tissue accumulation, immunogenicity, and local toxicity, while increasing cost and patient burden.^37,39^ Taken together, there is a critical need for strategies to both localize and sustain gene editing at target tissues to allow for reliable gene editing without off-target effects.

As an alternative to systemic/local injections, *in situ* delivery via biomaterial scaffolds localizes editing to target tissues and can extend the duration of nanoparticle release. The sustained release of non-viral CRISPR nanoparticles is typically achieved by physically embedding nanoparticles inside hydrogels or non-specifically adsorbing nanoparticles onto nanofibers or 3D printed scaffolds.^41^ For example, Zhong *et al*. extended gene editing over 14 days by adsorbing CRISPR nanoparticles to polydopamine coated nanofibers within a hydrogel matrix.^42^ Similarly, Ho *et al*. prolonged nanoparticle release over three days through adsorption into a fibrous hydrogel for the treatment of leukemia.^43^ However, non-specific nanoparticle adsorption strategies are limited by nanoparticle aggregation, un-even loading, disruption of nanoparticle integrity, and uncontrolled release kinetics, all of which reduce overall editing rates.^42,43^ Our objective was therefore to covalently tether nanoparticles to a biomaterial substrate to provide a defined biorthogonal conjugation system with spatiotemporal control over CRISPR gene editing.

To achieve the covalent tethering of non-viral nanoparticles, we identified the peptide-based SnoopTag/SpyTag bioconjugation system which enables rapid and irreversible covalent binding to complementary SnoopCatcher/SpyCatcher proteins.^44^ The SpyTag/SpyCatcher system is well-established for incorporating cell-specific ligands for targeted delivery of virus-like particles (VLPs), capsid-like particles, AAV, and nanoparticles, specifically for cancer immunotherapy.^45–52^ However, to the author’s knowledge these systems are yet to be applied to the covalent tethering of nanoparticles for nucleic acid delivery and gene editing applications. Therefore, we designed a SnoopTag functionalized RALA peptide and integrated it into RALA nanoparticles to enable covalent tethering to biomaterial substrates via a SnoopCatcher-SpyCatcher (Catcher) fusion protein.

Because RALA’s amphiphilic structure is essential for nanoparticle integrity, cell penetration, and endosomal escape, we bound SnoopTag-functionalized nanoparticles to zwitterionic microgels to avoid disrupting these properties.^25^ Zwitterionic materials contain balanced cationic and anionic groups within their molecular structure, creating ion-dipole interactions that form a hydration shell around the material to reduce nonspecific protein adsorption and biofouling.^53^ Furthermore, three-dimensional zwitterionic matrices are known to maintain the bioactivity of entrapped proteins and growth factors while supporting the proliferation and activity of seeded cells.^54,55^ Wheeler *et al*. recently demonstrated that MSCs cultured in annealed zwitterionic sulfobetaine methacrylate (SBMA) microgels increased extracellular matrix deposition due to a reduction in non-specific protein binding.^56^ Therefore, we adapted this system by functionalizing these zwitterionic microgels with the SpyTag peptide for covalently tethering RALA nanoparticles via the Catcher protein. We hypothesized that RALA-loaded zwitterionic microgels will provide an ideal substrate for CRISPR-delivery in a range of target tissues by enabling efficient RALA nanoparticle binding, controlled nanoparticle release rates, and sustained CRISPR action.

We demonstrate that SnoopTag-functionalized RALA efficiently integrates into RALA nanoparticles without compromising nanoparticle stability, cell viability, or transfection rates. Furthermore, we confirmed the presence of the SnoopTag peptide on the nanoparticle surface and its availability for bioconjugation with the Catcher protein. We also established the extended release of RALA nanoparticles from these microgels and their ability to sustain mRNA delivery to human primary MSCs seeded within our gene-loaded microgels. Finally, as a proof of concept, we demonstrated the prolonged CRISPR activation of *PRG4*, an important chondrogenic gene, demonstrating the promise of this microgel system for CRISPR-based regenerative strategies. These findings represent a novel approach for spatiotemporal control over CRISPR delivery by covalently tethering non-viral nanoparticles to a biomaterial scaffold.

## 2. Materials and methods

### 2.1. Materials

The RALA (WEARLARALARALARHLARALARALRACEA) and Snoop-RALA (KLGSIEFIKVNKSGESGSGWEARLARALARALARHLARALARALRACEA) peptides were produced by chemical peptide synthesis at Genscript, USA and provided as a lyophilized powder at 96.6% purity via high-performance liquid chromatography. SpyTag (Acryl – GGGSGGAHIVMVDAYKPTK) and RGD (Acryl-GGGSGGRGDS) were synthesized using standard solid phase peptide synthesis protocols as previously reported.^57^ Synthesis was performed using standard Fmoc-protected amino acids (Chemscene) on a Rink amide resin (Supra Sciences) unless otherwise noted. All amide couplings were performed in dimethylformamide (DMF) using the amino acid or acrylic acid, O-(6-chlorobenzotriazol-1-yl)- N,N,N′,N′-tetramethyluronium hexafluorophosphate (HCTU), and N,N-Diisopropylethylamine (DIPEA) at a 4:4:6 molar ratio, respectively. After each coupling, FMOC protecting groups were deprotected using 20% piperidine in dimethylformamide (DMF). A ninhydrin test was performed after each step to test for the presence of free amines and coupling or deprotection was repeated as necessary.

All peptides were cleaved using 95% trifluoroacetic acid (TFA), 2.5 % H_2_O, 2.5 % triisopropylsilane (TIPS) for 2–3 h at room temperature using 5 mL of cleavage solution per mM of peptide. The peptides were then precipitated and washed three times in diethyl ether by centrifuging at 5 minutes at 4,000 g. Peptides were purified using reverse-phase preparative HPLC (Agilent 218 Prep HPLC, Agilent Technologies, Santa Clara, CA, USA) on an Agilent 5 Prep-C18 column (150 mm × 21.2 mm, 5 μm pore size, 100 Å particle size). The purified peptide was confirmed using electrospray ionization mass spectrometry (ESI-MS; Applied Biosystems 3200 Q Trap, Foster City, CA, USA).

EZ Cap™ Cy5 EGFP mRNA (5-moUTP) was purchased from ApexBio. Plasmids for nanoluciferase (nanoluc) and CRISPRa mRNA production were purchased from Addgene. dSpCas9-VPR (Addgene plasmid # 205247; http://n2t.net/addgene:205247; <u>RRID:Addgene_205247</u>) was a gift from Rasmus Bak. pGEM4Z- T7-5’UTR-NanoLuc-3’UTR-A64 was a gift from Christopher Grigsby & Molly Stevens (Addgene plasmid # 203350 ; http://n2t.net/addgene:203350 ; RRID:Addgene_203350). dSpCas9-VPR and pGEM4Z-T7-5’UTR-NanoLuc-3’UTR-A64 plasmids were linearized using qPCR and purified using the GeneJET Gel Purification kit (Thermo Fisher). *In vitro* transcription was performed using the HiScribe T7 High-Yield RNA Synthesis Kit (NEB) with 3:1 N1- Methyl-pseudouridine 5’-triphosphate (Trilink Biotechnologies): unmodified UTP. mRNA was capped co-transcriptionally using the CleancapAG reagent (Trilink Biotechnologies) for dCas9- VPR and the 3′-O-Me-m^7^G(5’)ppp(5’)G RNA Cap Structure Analog (NEB) for nanoluc mRNA. mRNA was purified using the Monarch RNA Cleanup Kit (NEB). Yield and quality were assessed using a NanoDrop 2000 spectrophotometer (Thermo Scientific) and gel electrophoresis respectively.

### 2.2. Circular dichroism

CD was performed using a Jasco J-815 spectropolarimeter (Jasco International). Peptides were diluted to 0.2 mg/mL in a 20 mM NaH_2_PO_4_ 300 mM NaCl solution and loaded onto a 0.01 cm cuvette. Samples were recorded with a stepwise scanning mode standard sensitivity, 1 second digital integration time (D.I.T.), 1 nm bandwidth and data pitch, wavelengths of 280 – 190 nm and the ellipticity was calculated using equation 1.

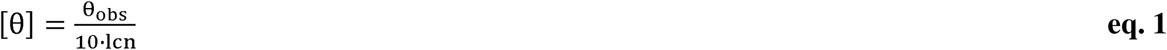

where: θ = mean residue molar ellipticity, θ_obs_= observed ellipticity in millidegrees, l = pathlength of cuvette (cm), c = molar concentration of the peptide, and n = number of amino acid residues.

### 2.3. Nanoparticle formation

Nanoparticles were formed using a modified protocol from previously reports.^14,24^ Briefly, RALA and snoop-RALA were resuspended at 1 mg/mL in nuclease free water and combined at various molar ratios. Nanoparticle formation per 1 μg of total RNA was performed by diluting RNA in water then adding 2.73 nmol of the combined peptide mixture for a final reaction volume of 25 μL. The mixture was incubated for 30 min at room temperature to allow for formation. Reactions were scaled accordingly for each experiment. For gene activations, 0.89 μg of dCas9 VPR mRNA and 0.11 μg of either non-targeting (control) or *PRG4* gRNAs were combined per 1 μg of total RNA in each nanoparticle reaction. Control non-targeting gRNAs were identified previously^58^ and *PRG4-*targeting gRNAs were optimized **(Fig. S1)**, all sequences are provided in **Table S1**.

### 2.4. Dynamic light scattering

The mean hydrodynamic diameter (size), zeta potential (surface charge), and polydispersity index (PDI) of nanoparticles were characterized using dynamic light scattering (DLS) (Zetasizer nano ZS, Malvern Panalytical, UK). After complexing, 50 µL of nanoparticle mixture was diluted 1:20 vol/vol in ultrapure water and loaded into polystyrene cuvettes or folded capillary zeta cells for size and charge using automated settings.^24^

### 2.5. Cell culture and monolayer transfections

Human bone marrow-derived mesenchymal stem cells (hMSCs) were obtained from RoosterBio (donor 310272). Cells were grown in expansion media (DMEM supplemented with 10% FBS (Gemini Bio) 1% penicillin/streptomycin (P/S, Thermo Fisher) for 7 days with media changes every 2–3 days. To passage cells, they were first treated with trypsin (0.25% trypsin, 0.1% EDTA in HBSS (Corning) for 5 minutes. Trypsin was then inactivated with expansion media. Cells were then pelleted by centrifugation for 5 minutes at 200 × g, resuspended in fresh expansion media and reseeded at 5.7 × 10^3^ cells cm^−2^ on tissue culture plastic treated for cell adhesion.

For transfection, MSCs were lifted using trypsin, diluted in 2x volume of expansion media, then centrifuged at 300 x *g* for 5 min. The cells were washed in optiMEM (Gibco) and finally resuspended at 1x10^5^ cells/mL. The cell suspension (0.1 mL cm^-2^) was then mixed with nanoparticles containing 0.15 μg cm^-2^ of RNA and incubated for 15 min at 37°C before plating in a 6 well-plate or 96 well-plate. After 6 h, the optiMEM was replaced with fresh expansion media.

### 2.6. Bioluminescent quantification

Nanoluc-transfected cells were treated with a final concentration of 10 mM furimizine (Selleck Chemical LLC). After 10 min, the well plates were imaged using a Perkin Elmer Lumina III IVIS. ROIs were drawn around each well to quantify the average radiance (p s^−1^ cm^−2^ sr^−1^) of each sample.

### 2.7. Flow cytometry

Samples were prepared for flow cytometry by detaching cells with trypsin then resuspending in PBS. Samples were assessed using a Cytoflex flow cytometer (Beckman Coulter, USA). Samples were recorded at a variable flow rate to maintain the event rate at 400 events s^−1^. The FSC, SSC, FITC, and APC channel gains were set to 20, 20, 1, and 25, respectively. The resulting data were gated on FSC-A and SSC-A to remove debris then gated for FSC-A and FSC-H to select single cells. Fluorescent thresholds were gated using non-transfected cells as reference and the transfection rate was defined as the percentage of positive cells compared to the total number of single cells. The cell viability was defined as the percentage of total single cells in transfected samples versus a non-transfected control. The transfection yield was based on previously established metrics.^24,59,60^ Yield was defined as the percentage of GFP compared to the total single cell count of the non-transfected control.

### 2.8. Gene expression analysis

Total RNA was extracted using the Trizol reagent (Invitrogen), purified using PureLink RNA Mini Kit (Invitrogen), and reverse transcribed using SuperScript IV VILO Master Mix (Invitrogen). Quantitative PCR (qPCR) was performed using TaqMan Fast Advanced Master Mix for qPCR (Applied Biosystems) and specific gene expression assays (Applied Biosystems). TaqMan gene expression assay primers were added in multiplex with the *RPL13A* (Hs04194366_g1) housekeeping gene primer labelled with VIC and *PRG4* (Hs00981633_m1) labelled with FAM. Reactions were prepared according to the manufacturers protocol in a 0.2 mL MicroAmp optical 96-well reaction plate (Applied Biosystems) and PCR was performed in a QuantStudio3 real-time PCR system (Applied Biosystems, USA) with the ΔΔCt method. All fold changes were quantified relative to a non-transfected control and a control transfected with gRNA targeting the *AAVS1* safe harbor locus as a non-targeting control.

### 2.9. Microgel synthesis

Zwitterionic microgels were produced as previously reported.^56^ Briefly, [2-(methacryloyloxy) ethyl] dimethyl-(3-sulfopropyl) ammonium hydroxide] (SBMA, Sigma–Aldrich) was resuspended at a 10% w/v concentration and 5% w/v of poly(ethylene glycol) dimethacrylate (PEGDMA, MW 750, Sigma-Aldrich) in HEPES buffer with 0.4% of photoinitiator 2,2′- Azobis[2-methyl-N-(2-hydroxyethyl) propionamide] (VA-086, FUJIFILM).

Microgels were fabricated using an emulsion method in a polydimethylsiloxane (PDMS) microfluidic device channel height of 35 µm and reservoir height of 250 µm.^56,61,62^ The continuous oil phase consisted of Novec 7500 oil (3 M) and 0.75 wt.% Picosurf (Sphere Fluidics) and the aqueous phase consisted of 10% w/v polymer. Each solution was added to their respective syringes with a 21G needle connected via tubing. The continuous phase flow rate (80 µL min^−1^) was double that of the dispersed phase (40 µL min^−1^) using syringe pumps (NE- 1000 and KD Scientific). Microgels were collected in a glass vial containing 10 mL of mineral oil, exposed to 20 mW cm^-2^ for 3 min to crosslink, then incubated overnight to allow swelling.

After removal of mineral oil and Novec 7500 oil, a solution of 20 wt.% 1H,1H,2H,2H-Perfluoro- 1-octanol (PFO, Sigma) diluted in Novec 7500 oil was added to an equal volume of microgels. To disperse the microgels, 25 mM HEPES buffer at pH 7.4 (HEPES) was added and centrifuged at 2000 x *g* for 5 min. The bottom layer of oil and top layer of HEPES was removed and washed in HEPES 3x in 2000 x *g* for 5 min. For sterile experiments, microgels sat in 70% isopropyl alcohol (IPA) for 15 min, washed 3x with 70% IPA, and then washed 3x with sterile HEPES buffer with 1% P/S.

To functionalize the microgels, microgels were pelleted by centrifugation at 3000 x g for 5 min, then resuspended in an equal volume of 1.6 mM Acryl-SpyTag, 3.2 mM Acryl-RGD, and 0.5% LAP. The microgels were then treated with 405 nm UV light (25 mW cm^-2^) for 3 min with regular mixing. The microgels were finally washed 3 times with 10x volume of HEPES to remove unbound peptides.

### 2.10. Protein synthesis

Plasmids for protein production were purchased from Addgene. pET28a SpyCatcher-SnoopCatcher was a gift from Mark Howarth (Addgene plasmid # 72324; http://n2t.net/addgene:72324 ; RRID:Addgene_72324).^44^ pET28a-SnoopTag2-sfGFP was a gift from Mark Howarth (Addgene plasmid # 201810).^63^ SnoopCatcher-SpyCatcher and SnoopTag- GFP pDNA were cultured in LB Broth supplemented with 100 μg/mL carbenicillin and 50 μg/mL kanamycin, respectively, then purified using the GeneJET Plasmid Miniprep Kit (Thermo Fisher). The plasmids were then transformed into BL21-codon plus RIPL cells (Agilent) using heat shock at 42°C for 20 min, and single colonies were selected from LB-agar plates treated with the necessary antibiotics.

Single colonies were grown in 10 mL LB medium for 16 h at 37°C with shaking at 200 rpm. Starter cultures were diluted 1:100 into 1 L LB supplemented with 0.8% (w/v) glucose and either 50 μg/mL kanamycin or 100 μg/mL ampicillin (8 g glucose and 100 mg ampicillin or 50 mg kanamycin per liter of LB). Cultures were grown at 37°C with shaking at 200 rpm until reaching an OD of 0.5 (approximately 2.5–3 h). Protein expression was induced with 0.42 mM isopropyl β- d-1-thiogalactopyranoside (IPTG). After induction, Catcher cultures were grown 4 h at 30°C and 200 rpm, and SnoopTag-GFP cultures were incubated for 18 h at 22°C and 200 rpm. Cells were pelleted by centrifugation at 5,000 × *g* for 10 min in a pre-weighed 500 mL ultracentrifuge tube, and the wet cell pellet mass was recorded. Pellets were stored at −20 °C to −80°C overnight.

Cell pellets were resuspended in 5 mL of B-PER Complete Reagent (Thermo Fisher) per gram of biomass and incubated for 15 min at room temperature with gentle agitation. The cell lysates were then transferred to ultracentrifuge tubes and clarified by centrifugation at 16,000 x *g* for 20 min.

Proteins were purified using the HisPur Ni-NTA Spin Purification Kit (Thermo Fisher) using 20 mM Sodium Phosphate 300 mM Sodium Chloride buffer with imidazole concentrations and wash steps listed in table 1.

**Table 1.** Wash steps for Immobilized Metal Affinity Chromatography (IMAC).

| Step | [Imidazole] (mM) | Volume (mL) | Incubation time (min) |
| --- | --- | --- | --- |
| Column equilibration | 10 | 6 | 2 |
| Column loading | 5 | 15 | 30 |
| Wash I | 25 | 6 | 2 |
| Wash II & III | 50 | 6 | 2 |
| Elution I | 250 | 3 | 5 |
| Elution II | 500 | 3 | 5 |

The eluate was dialyzed into 20 mM Tris·HCl, pH 8, overnight before further purification via anion exchange chromatography (AEX). AEX was performed using a 1 mL HiTrap Q HP AEX column (Cytivia) on a ÄKTAprime plus instrument (Cytiva). Proteins were eluted using a 20 mL gradient from 0-500 mM NaCl with a 0.5 mL/min flow rate and 0.25 mL fractions collected. Finally, the eluted proteins were dialyzed into 25 mM Tris, 2.7 mM KCl, and 137 mM NaCl for the Catcher proteins or 137 mM NaCl, 2.7 mM KCl, 10 mM Na2HPO4, 2 mM KH2PO4, pH 7.4 for SnoopTag-GFP. Proteins concentrations were determined by A280 readings on the Nanodrop2000, and protein purity was assessed by denaturing SDS PAGE.

### 2.11. Denaturing SDS PAGE

1 μg of protein was combined with 4x NuPAGE LDS Sample Buffer (Thermo Fisher) and 50 mM Dithiothreitol (DTT) then incubated at 75°C for 10 min. Samples were run at 200 V for 35 min in NuPage Bis-Tris Mini Protein Gels, 4–12%, 1.0 mm using 1x NuPAGE MES running buffer with the inner chamber supplemented with NuPAGE antioxidant. Gels were stained using the SimplyBlue SafeStain (Thermo Fisher) and imaged using the Odyssey FC gel imager (Licor).

### 2.12. Non-denaturing PAGE

Samples were combined with 4X NativePAGE Sample Buffer (Thermo Fisher) and diluted with DI water to a final volume of 10 μL. Samples were run at 150 V for 115 min in NativePAGE Novex Bis-Tris Mini Protein Gels using Dark Blue Cathode Buffer (20X NativePAGE Running Buffer and 20X Cathode Additive, Thermo Fisher) in the inner chamber and Anode Buffer (20X NativePAGE Running Buffer, Thermo Fisher) in the outer chamber. The gels were destained using 40% methanol and 10% acetic acid in ultrapure water with buffer exchanged every 30 min then imaged using the Odyssey FC gel imager (Licor).

### 2.13. Snoop-Catcher binding assay

Snoop-RALA nanoparticles were prepared and diluted with 250 mM HEPES for a final concentration of 25 mM HEPES, then the Catcher protein was added at a 10:1 SnoopTag:Catcher molar ratio. The nanoparticles and Catcher protein were incubated at room temperature with constant mixing for 90 min. Enough sample for 1 μg of Catcher protein was then analyzed by denaturing and non-denaturing SDS PAGE. Band intensities were quantified using the Gel Analyzer function in ImageJ.^64^

### 2.14. SnoopTag-GFP binding assay

Microgels were incubated with 10 μg of the catcher protein and 10 μg of SnoopTag-GFP. The gels were incubated on a spinning wheel for 2 h then washed 3 times with 10x volume of HEPES. Microgels were imaged on a Keyence BZX-800 microscope (Keyence Corporation, USA).

### 2.15. Nanoparticle binding to microgels

For each reaction, 50 μL of microgels were mixed with 7.5 ng of Catcher in 100 μL of HEPES and incubated for 1 h with constant agitation. The microgels were then centrifuged at 3000 x *g*, and the supernatant was aspirated to remove unbound Catcher protein. Snoop-RALA nanoparticles were prepared with 1.5 μg of total RNA per reaction, diluted with 2x volume of HEPES, then added to the microgels. This reaction was incubated at room temperature with constant mixing for 2 h. Finally, the microgels were centrifuged at 3,000 x *g*, and the supernatant was aspirated.

To visualize nanoparticle binding, Snoop-RALA nanoparticles made with Cy5 labelled mRNA were incubated with SpyTag functionalized microgels with or without the Catcher protein. The microgels were washed three times with 20x volume of HEPES buffer, with 10 min of constant agitation per wash. Microgels were imaged on a Keyence BZX-800 microscope using the Cy5 filter.

### 2.16. Nanoparticle release profile

Cy5 labelled nanoparticles were bound to SpyTag functionalized microgels and the supernatant after the binding reaction was saved. An equal volume of nanoparticles was saved as a control for cumulative release (nanoparticle control). The microgels were incubated in 300 μL of HEPES with constant agitation. On days 1, 3, 5, 7, 10, and 14, the microgels were centrifuged at 3,000 x *g* and the supernatant was aspirated, saved then replaced by fresh HEPES. After the last time point, 50 μL of 10% SDS was added to each sample or the remaining microgels in 300 μL of HEPES and incubated for 1 h at room temperature. The fluorescence of each sample at each time point was measured in triplicate at 640/680 excitation/emission using a SpectraMax iD3 plate reader. The fluorescent intensity of each sample was normalized by subtracting the intensity of 1.75% SDS in HEPES. Cumulative release was calculated by summing the average fluorescence intensity across three replicates for each sample and reported as the percentage of the nanoparticle control.

### 2.17. Cell seeding on microgels

Cells were lifted using trypsin, washed with optiMEM, counted, and resuspended in optiMEM at 2 x 10^6^ cells/mL. For each reaction, 100 μL of cell solution was mixed gently with 50 μL of nanoparticle-bound microgels by inverting the tube, then the mixture was pipetted into a single well of a 96 well-plate for a final concentration of 200,000 cells/well. The well plates were then centrifuged at 300 x *g* for 2 min on each side to jam the microgels and 100 μL of optiMEM was added on top of the jammed scaffold. Cell-seeded scaffolds were analyzed using IVIS, fluorescent imaging, or qPCR as described at the corresponding time points. Images of GFP expression within the microgel scaffolds were taken as z-stacks containing 90-150 images with 2.8 μm spacing between each image. Images were 3D projected using the brightest point projection method with interpolation in imageJ.^64^

### 2.18. Statistics

Statistical analyses were performed using GraphPad Prism (version 11.0.0). For groups compared by statistical analysis, data were tested for normality using the Shapiro–Wilk test. For normal data with three or more groups, a one-way ANOVA was used with a Brown–Forsythe test and a Tukey post-test. Non-normal data were analyzed using the Kruskal–Wallis test with multiple comparisons via Dunn’s test. Multi-day experiments were analyzed by two-way ANOVA with Tukey post-test. Numerical and graphical results were displayed as mean ± standard deviation (SD). Significance was accepted at *p* < 0.05. Sample size (*n*) is indicated within the corresponding figure legends. Each datapoint represents an individual sample from a single experiment.

## 3. Results

### 3.1. Non-viral delivery of mRNA in monolayer culture results in transient gene expression, limiting the duration of CRISPR activity

To characterize the temporal dynamics of mRNA-mediated transgene expression, mRNAs encoding GFP, nanoluciferase (nanoluc), or dCas9-VPR were delivered to MSCs via RALA nanoparticles, and transgene activity was monitored over time (**Fig. 1A**). Nanoluc expression, assessed by IVIS bioluminescence imaging over 21 days, remained strong for the first three days before declining sharply for the remainder of the observation period (**Fig. 1B**). Consistent with this trend, the transfection of GFP mRNA results in the rapid decrease of GFP mean fluorescence intensity (MFI) over a 5-day period as confirmed by fluorescent imaging (**Fig. 1C**) and single-cell analysis of GFP expression by flow cytometry (**Fig. 1D**). To determine the functional impact of this transient expression on CRISPR activity, we delivered dCas9-VPR and gRNA targeting the *PRG4* gene, a gene previously identified in our lab as a promising CRISPRa target for MSC chondrogenesis through *in silico* CRISPR screens.^65^ *PRG4* encodes lubricin, a proteoglycan that lubricates the superficial layer of articular cartilage and serves as a key marker of chondroprogenitor cells and MSC chondrogenesis.^65–67^ Notably, unlike genetic editing tools that permanently alter DNA sequence, CRISPRa does not modify the genome, so activation persists only as long as the delivered mRNA is expressed. Concurrent with nanoluc and GFP data, *PRG4* activation peaked at day 3, but no activation was observed by day 7, indicating that CRISPR activity following mRNA delivery is sustained for approximately 3-5 days (**Fig. 1E**). These results highlight the need for delivery systems capable of sustaining and localizing CRISPR activity over extended time frames, particularly for regenerative medicine applications such as cartilage repair, where chondrogenic differentiation and tissue repair unfold over weeks.

**Figure 1.**
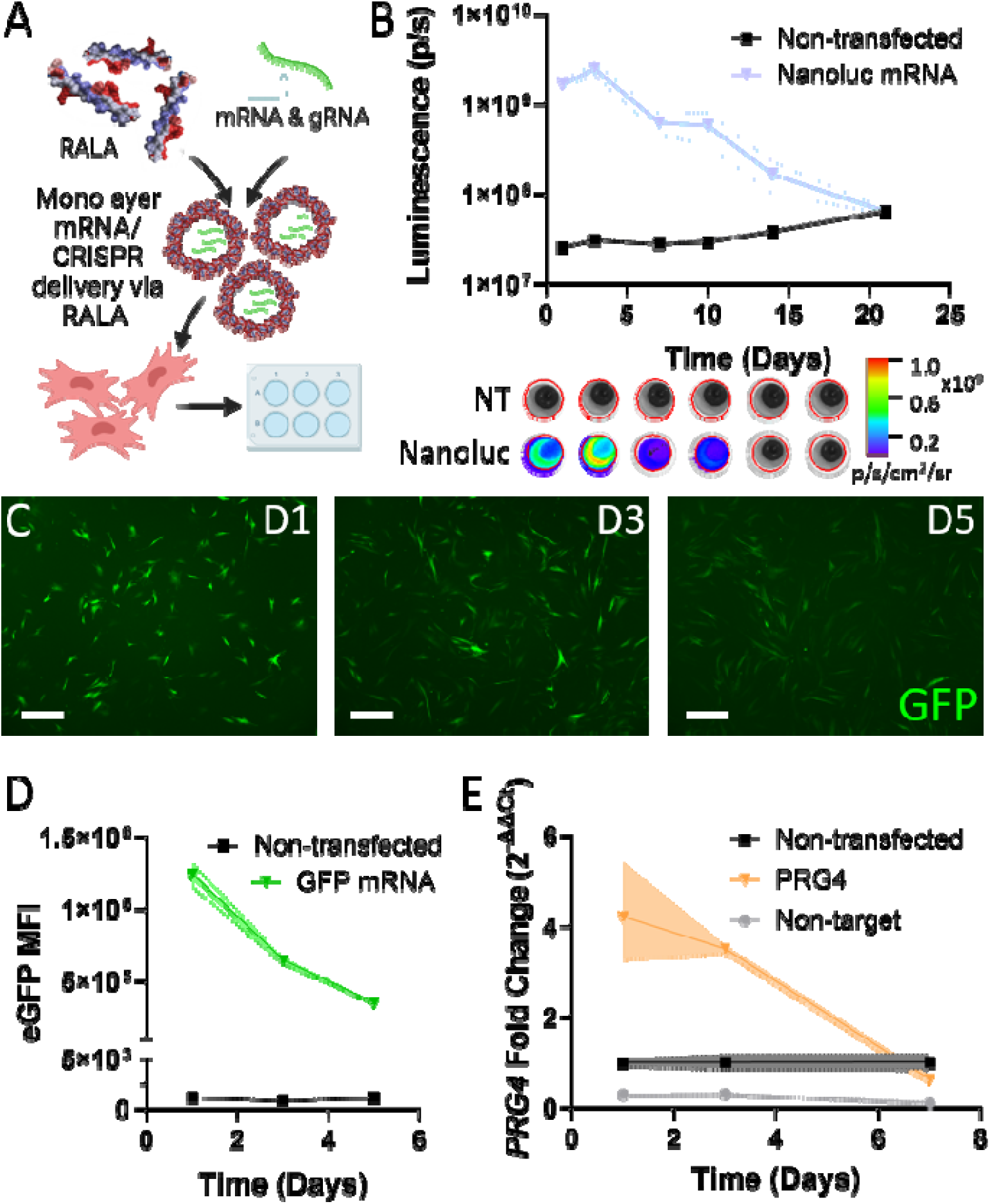
Transient gene expression following non-viral RALA-based mRNA delivery. **A)** The temporal expression profiles of RALA-transfected mRNAs in monolayers were analyzed. **B)** The luminescence of MSCs transfected with nanoluc mRNA was characterized by IVIS ROI quantification over 21 days with representative images of the luminescent flux (p/s/cm^2^/sr) at each time point (n=3). RALA was then used to transfect GFP mRNA which was monitored by **C)** fluorescent imaging and **D)** flow cytometry analysis (n=3). Data are plotted as the mean with shading representing standard deviation, scale = 500μm. **E)** MSCs were then transfected with dCas9-VPR mRNA and *PRG4* gRNA and analyzed by RT-qPCR on days 1, 3, and 7. Data are plotted as geometric mean ± geometric standard deviation of the fold change (2^-ΔΔCt^) relative to non-transfected controls (black lines) and the *RPL13A* housekeeping gene (n=3). Gene activation is also compared to a non-targeting control (gray lines).

### 3.2. Engineering SnoopTag functionalized RALA nanoparticles

To improve the spatial and temporal release kinetics of RALA-based cell delivery, we identified the SnoopTag/SpyTag system to tether RALA nanoparticles to zwitterionic microgels.^44^ SnoopTag, which covalently binds with the SnoopCatcher protein, was added to the N-terminus of the RALA peptide separated by a SGESGSG linker to make the Snoop-RALA peptide. Since the alpha helicity of RALA is essential for nanoparticle complexing, cellular delivery, and endosomal escape, we used AlphaFold 3 to compare the predicted secondary structure of Snoop- RALA to the RALA peptide.^25,68^ The resulting predictions indicate that Snoop-RALA maintains the alpha helicity of its RALA domain with the SnoopTag free on the N-terminus. (**Fig. 2A**). Circular dichroism further confirms that Snoop-RALA maintains this partial alpha-helical structure (**Fig. 2B**). To form SnoopTag functionalized nanoparticles, Snoop-RALA and RALA were mixed at various molar ratios before adding to mRNA (**Fig. 2C**). DLS was used to determine the influence of Snoop-RALA on the size and charge of the resulting nanoparticles. We demonstrate that nanoparticle size increases slightly with the addition of SnoopTag-RALA (**Fig. 2D**). Nanoparticle charge increased with higher percentages of Snoop-RALA up to 20%, consistent with Snoop-RALA’s higher net charge (+6) relative to unmodified RALA (+5). However, nanoparticle charge decreased above 20% Snoop-RALA, likely due to instability (**Fig. 2D**). These trends were further confirmed by the transfection of MSCs. The percentage of transfected cells and cell viability had an inverse relationship as the Snoop-RALA percentage increases (Fig. S2). This relationship led to a consistent transfected cell yield and mean fluorescent intensity with up to 20% Snoop-RALA (**Fig. 2E**, Fig. S1C). The reduction in transfection rate and improved viability can be explained by the increasing surface coverage by SnoopTag, which reduces interactions with the cell membrane as well as non-specific nanoparticle adhesion, allowing for improved cell adhesion.^69^ Together these results indicate that up to 20% Snoop-RALA can be incorporated into the nanoparticles without significantly impacting nanoparticle formation and cell transfection.

**Figure 2.**
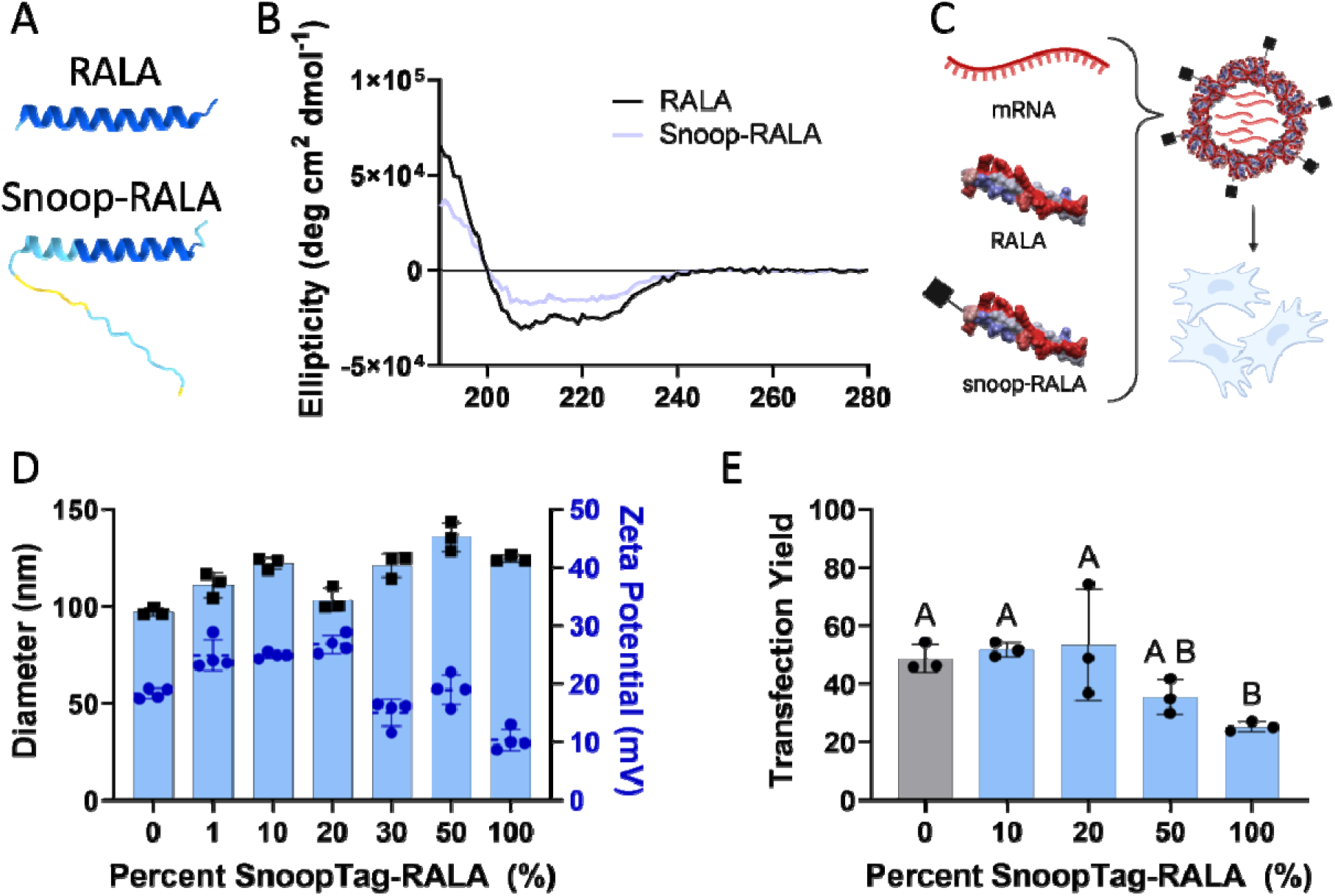
Design and characterization of SnoopTag-functionalized RALA nanoparticles. The secondary structures of the Snoop-RALA peptide were characterized via **A)** AlphaFold 3 predictions and **B)** circular dichroism indicating that the RALA domain maintains its essential alpha-helical conformation. **C)** Nanoparticles were then formed by mixing different molar percentages of RALA and Snoop-RALA with mRNA. The resulting nanoparticles were then characterized by **D)** DLS to determine their size (n=3) and charge (n=4) and **E)** their transfection yield (percent transfected normalized with viability) (n=3) to determine the overall nanoparticle functionality. DLS and transfection data are plotted as the mean ± standard deviation. Statistically similar groups are denoted with the same letters (one-way ANOVA, p < 0.05).

### 3.3. SnoopCatcher-SpyCatcher efficiently binds Snoop-RALA nanoparticles and SpyTag functionalized microgels

After successfully integrating SnoopTag into RALA nanoparticles, we sought to confirm the efficacy of the SnoopTag/SpyTag bioconjugation systems for tethering the nanoparticles to zwitterionic microgels. First, we evaluated the availability of the SnoopTag residues on the surface of Snoop-RALA nanoparticles. Nanoparticles were formed with 20% Snoop-RALA then reacted with the Catcher protein at a 10:1 Snoop-RALA:Catcher molar ratio. The resulting conjugate was analyzed by non-denaturing and denaturing PAGE (**Fig. 3A**). In denaturing PAGE, SDS disrupts the nanoparticles, allowing the bound snoop-RALA to run through the gel. Gel analysis demonstrates that 100% of the Catcher is shifted higher in the gel, indicating that it is bound to Snoop-RALA (**Fig. 3B**). Alternatively, the absence of SDS in non-denaturing PAGE maintains the integrity of the nanoparticles which are much larger than the mesh size of the polyacrylamide. The absence of a Catcher band in the non-denaturing gel confirms that the Catcher protein is integrated into the surface of the nanoparticles (**Fig. 3A-B**). Next, we confirmed the binding of the Catcher protein to the SpyTag-functionalized microgels. SpyTag microgels were reacted with SnoopTag-GFP in the presence or absence of the Catcher protein and analyzed by fluorescent microscopy. A strong fluorescent intensity was observed only when the catcher protein was added to the reaction indicating that the Catcher protein efficiently tethers proteins to the surface of zwitterionic microgels (**Fig. 3C**).

**Figure 3.**
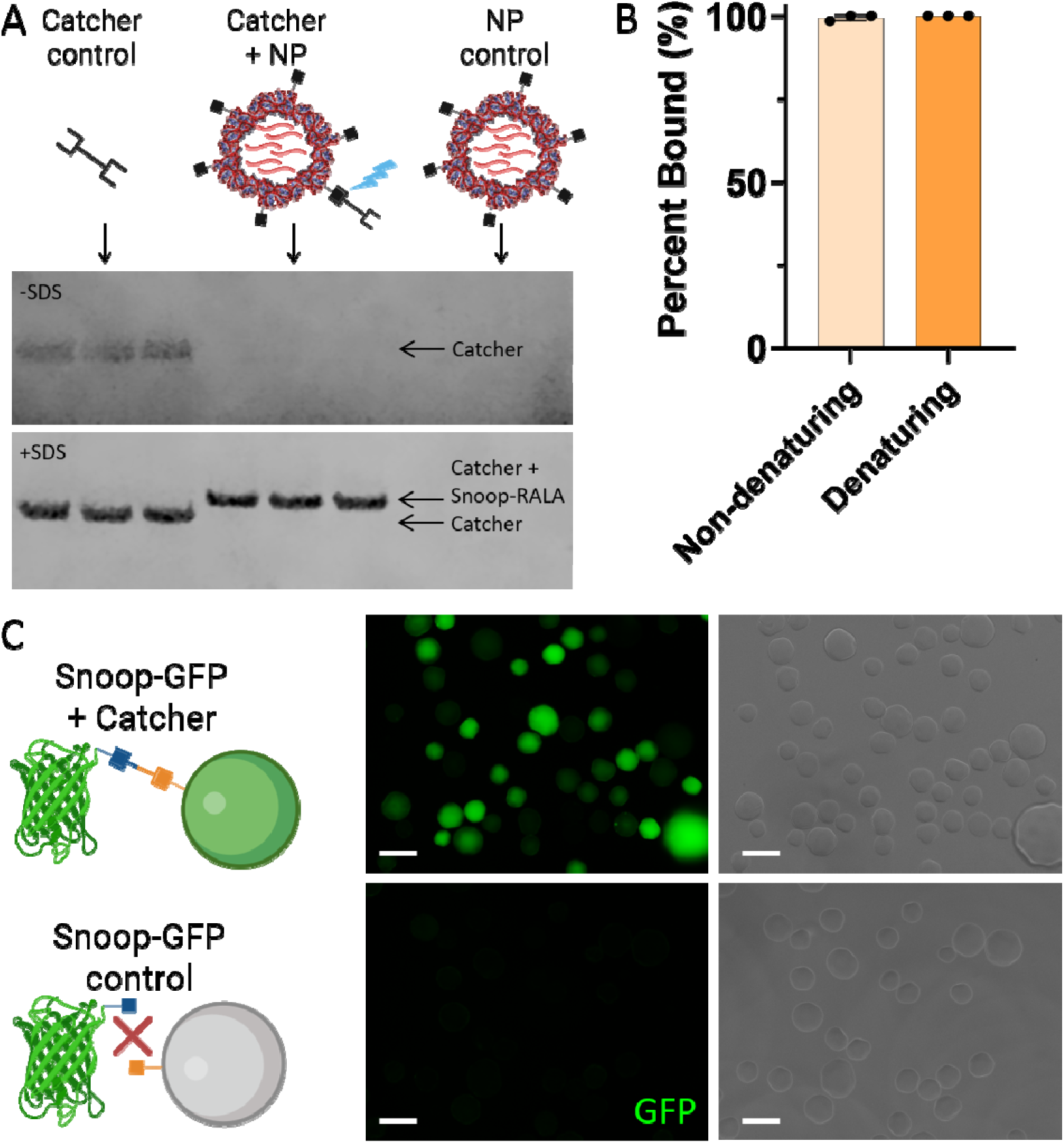
Validation of bioconjugation reaction validation. A) SnoopTag-RALA nanoparticle were reacted with the SnoopCatcher-SpyCatcher (Catcher) protein then analyzed by non-denaturing (- SDS) or denaturing (+ SDS) PAGE. B) Binding efficiencies as quantified by ImageJ analysis of PAGE band intensities (n=3). Plotted as the mean ± standard deviation. C) Catcher and SnoopTag binding to microgels is visualized by binding a SnoopTag-GFP fusion protein to the microgels, scale = 100 μm.

### 3.4. Covalent tethering to zwitterionic microgels results in sustained release of Snoop-RALA nanoparticles

After validating the Catcher system, we tethered Snoop-RALA nanoparticles to SpyTag- functionalized nanoparticles. First, we confirmed binding specificity by reacting Cy5-labelled nanoparticles with the microgels in the presence or absence of the Catcher protein (**Fig. 4A**). Again, we demonstrate a strong fluorescent signal only when the catcher protein is incorporated, confirming the Catcher protein is covalently linking the nanoparticles and microgels. Furthermore, this specificity also demonstrates the importance of the zwitterionic properties for preventing non-specific adsorption and nanoparticle aggregation. Next, we determined the release kinetics of nanoparticles bound to microgels. Cy5 labelled nanoparticles were reacted with the Catcher protein and SpyTag-functionalized microgels and incubated in HEPES buffer with constant agitation for 14 days with the buffer collected at regular intervals (**Fig. 4B**). The nanoparticles released slowly, plateauing at 21.5 ± 3.6 % released on day 14 (**Fig. 4B).** The remaining microgels were then treated with SDS to release the remaining Cy5 labelled mRNA, confirming that 59.5 ± 8.7% of nanoparticles remained bound to the microgels (**Fig. 4B**). These results are further verified by fluorescent imaging with strong Cy5 signal observed through 14 days (**Fig. 4C**).

**Figure 4.**
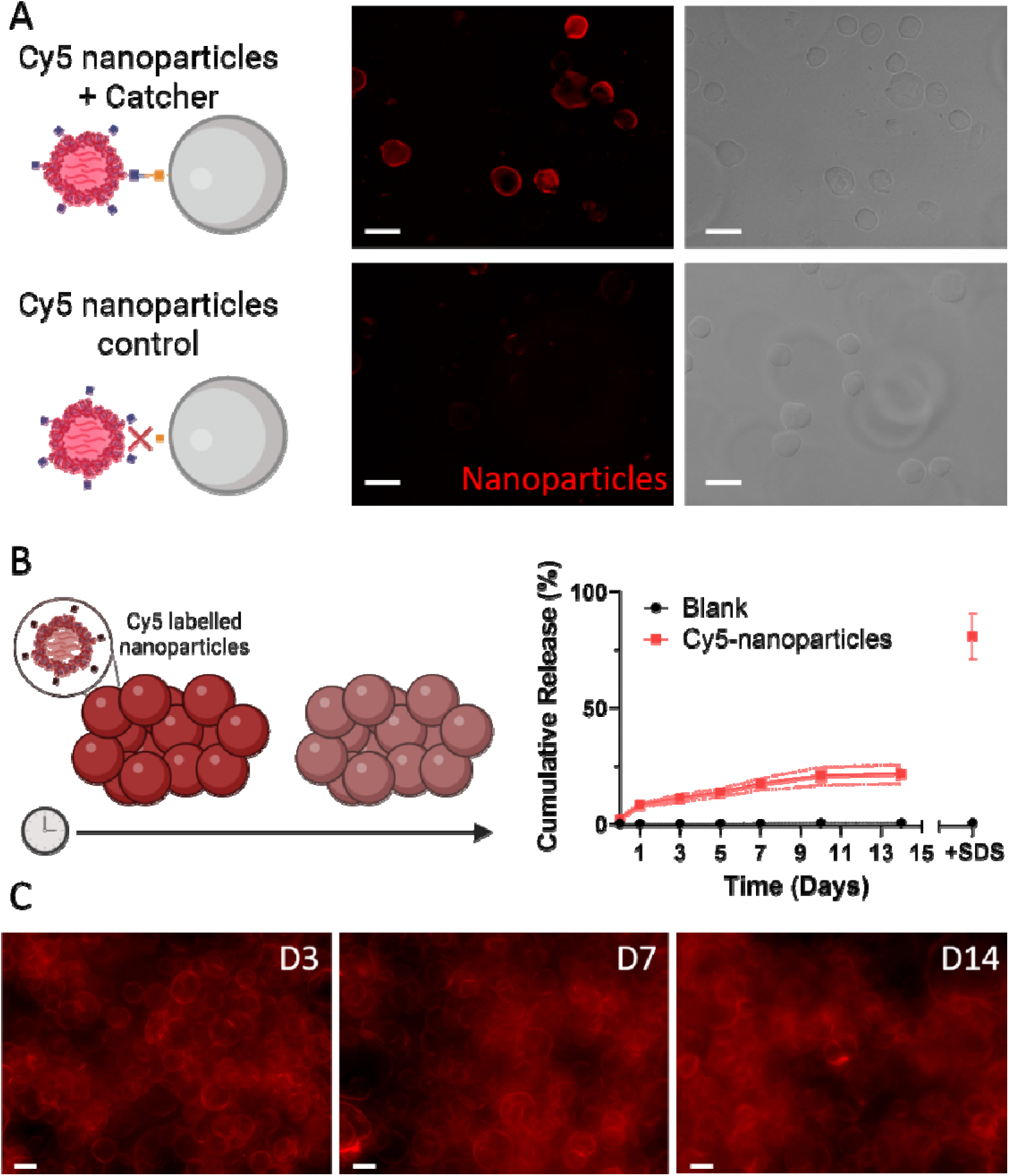
Snoop-RALA nanoparticles bind specifically to zwitterionic microgels for sustained nanoparticle release. **(A)** Snoop-RALA nanoparticles complexed with Cy5-labeled mRNA were reacted with SpyTag-functionalized microgels in the presence or absence of Catcher protein, demonstrating Catcher-dependent binding of nanoparticles to the microgels. Scale = 250 μm. **B)** The cumulative release rate of nanoparticles from Catcher-bound microgels was quantified over 14 days (n=3) and plotted as the mean ± standard deviation. **(C)** Nanoparticle retention was also visualized by representative images, scale = 100 μm.

### 3.5 Snoop-RALA loaded microgels sustain mRNA delivery and CRISPR gene editing in seeded human MSCs

With our system optimized, we sought to characterize the transfection of primary human MSCs within a 3D matrix of Snoop-RALA loaded microgels (**Fig. 5A**). We first tethered eGFP mRNA nanoparticles to SpyTag microgels, added cells, then jammed into a 3D porous scaffold. Across all timepoints assessed (days 3, 7, 10, and 14), GFP^+^ cells were observed in two morphologies, either contoured around the microgel surfaces or rounded within the void spaces. (**Fig. 5B**). Bright GFP expression was observed 14 days post cell seeding, indicating the sustained nanoparticle uptake and expression of GFP mRNA nanoparticles. Prolonged luciferase activity was also observed in microgels loaded with nanoparticles containing nanoluc mRNA (**Fig. 5C**). In both the presence and absence of the Catcher linker protein, luminescence peaked after 3 days, and this signal was maintained through day 10. After 10 days, the signal decreased slowly through day 21 but never fell below the signal recorded on day 1, indicating the sustained uptake of nanoluc mRNA (**Fig. 5C**). This sustained signal contrasted with monolayer transfection, where luminescence fell below day 1 levels within one week (**Fig. 1B**). Finally, Snoop-RALA nanoparticles encapsulating dCas9-VPR mRNA and either non-targeting or *PRG4*-targeting gRNA were loaded onto the microgels to assess CRISPRa activity compared to monolayer transfection (**Fig. 5D**). On day 3, *PRG4* was upregulated 71.0 ± 3.8-fold and 71.8 ± 1.6-fold with and without the Catcher protein linker, respectively (**Fig. 5D**). These fold-changes were much higher than gene activation in monolayer transfection which upregulated *PRG4* a maximum of 3.5 ± 0.1-fold (**Fig. 1E**). Interestingly, the non-targeting control upregulated *PRG4* 11.1 ± 1.9- fold in the microgel system versus 0.3 ± 0.01-fold in monolayer indicating unique cellular responses to transfection in 3D microgels vs 2D cell-culture plastic (**Fig. 5D**). Although gene activation was consistent with and without the Catcher-protein tethering the nanoparticles to microgels on day 3, the absence of the Catcher protein led to increased CRISPRa activity on day 7 with an 18.4 ± 6.9-fold increase in *PRG4* expression relative to non-transfected controls versus 10.0 ± 2.7-fold with the Catcher protein linker (**Fig. 5D**). Still, both outperformed monolayer conditions which had a fold change of 0.6 ± 0.1-fold (**Fig. 1E**). This trend continued at day 14 with upregulated expression of *PRG4* in groups with and without catcher protein, although this upregulation was not significantly superior to the non-targeting control, highlighting the transcriptomic effects of the material substrate and delivery vector (**Fig. 5D**). These results indicate that in the presence and absence of the Catcher protein, nanoparticles are entrapped within the packed microgels and are available for transfecting MSCs, demonstrating a significant improvement of CRISPR temporal dynamics over standard monolayer protocols.

**Figure 5.**
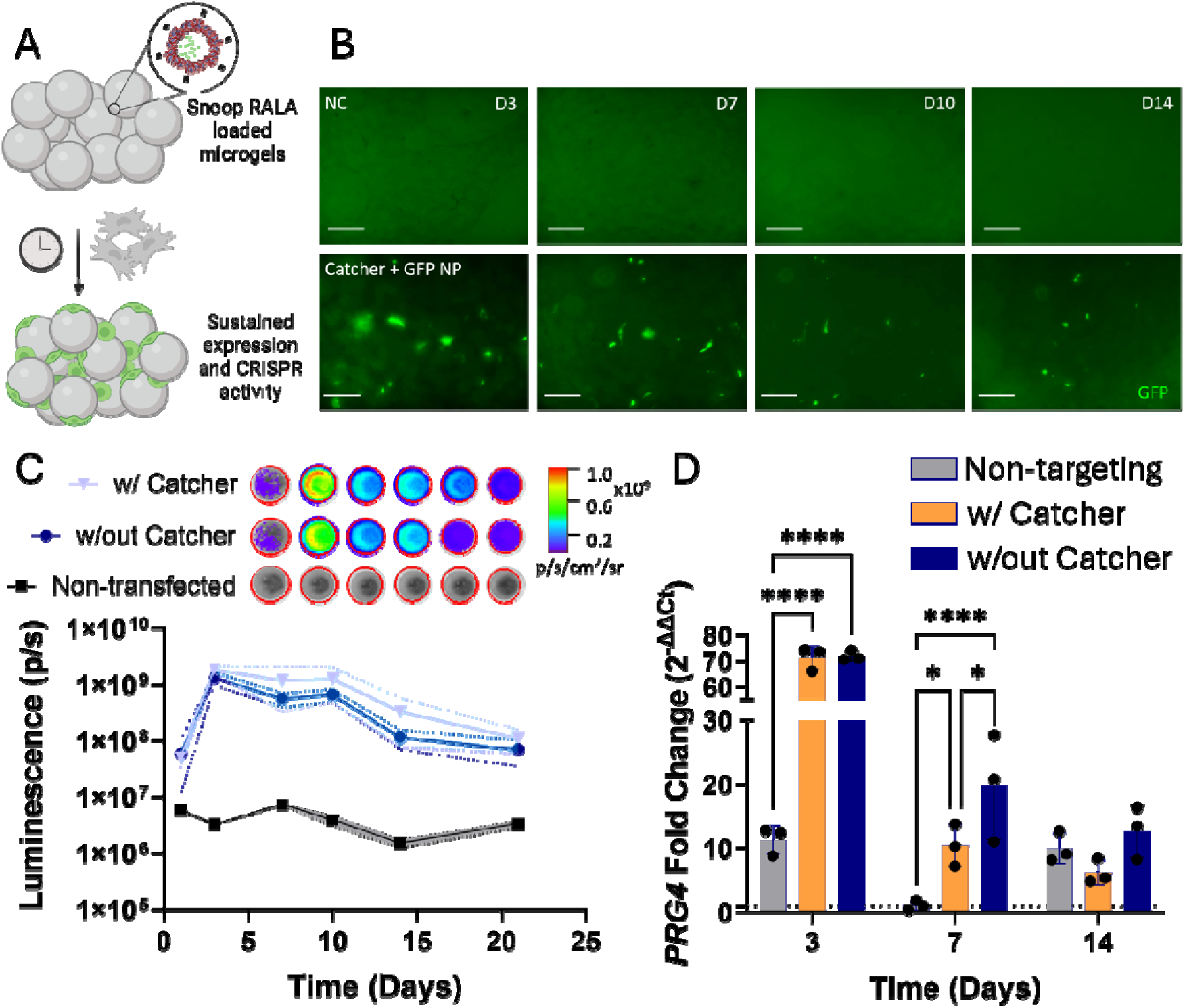
Sustained mRNA expression and CRISPR activity in Snoop-RALA loaded microgels. **A)** MSCs were seeded inside of jammed microgel scaffolds loaded with GFP, nanoluc, or dCas9- VPR mRNA. **B)** Representative fluorescent images of GFP expression within the microgels, scale bar = 250 μm. **C)** Luminescent intensity of MSCs seeded on nanoluc-mRNA loaded scaffolds with or without the Catcher protein linker compared to non-transfected controls with representative images of the luminescent flux (p/s/cm^2^/sr) included. Data is plotted as the mean with shading representing standard deviation. **D)** *PRG4* fold change relative to non-transfected controls in cells seeded on microgels loaded with nanoparticles containing dCas9-VPR mRNA and either non-targeting or *PRG4-*targeting gRNA. *PRG4* activation was compared with or without the Catcher protein tethering nanoparticles to the microgels. Plotted as the geometric mean ± geometric standard deviation. * denotes significance p < 0.05 (*n* = 3), \*\*\*\**p* < 0.0001 (*n* = 3).

## 4. Discussion

The effective delivery of CRISPR machinery to therapeutic cell types remains one of the largest barriers to clinical translation. Effective CRISPR therapies require efficient cellular delivery with minimal unintended transcriptomic effects that could mask the true effects of gene editing or induce inflammation. We have previously demonstrated that the RALA cell penetrating peptide delivers CRISPR mRNA and RNP for efficient gene editing with reduced inflammatory effects compared to gold standard non-viral systems such as lipid-based or polymeric nanoparticles.^14^ RALA has achieved efficient gene knock-in, knock-out, and transcriptional activation in primary MSCs and is a safe approach for *in vivo* gene delivery.^24–26,70–72^ Yet, the translation of RALA- based strategies is limited by poor biodistribution, with nanoparticles accumulating in the lungs and spleen, and transient gene expression which limits editing rates at target sites.^25^ Therefore, we sought to design a platform for the sustained release of RALA nanoparticles via covalent linkage to 3D biomaterial substrates. We identified the SnoopTag/SpyTag bioconjugation systems that rapidly and irreversibly bind to their complementary SnoopCatcher/SpyCatcher proteins in aqueous solutions at neutral pH with simple procedures, making them ideal for preserving the integrity of RALA nanoparticles.^44,56,73^ Furthermore, the SpyTag/SpyCatcher system is well established for modifying drug delivery systems, although, to the author’s knowledge, this is the first application of this or any bioconjugation system for covalent binding of nanoparticles to biomaterial scaffolds.^49–52,74^

To integrate these systems, we first designed Snoop-RALA, a RALA and SnoopTag fusion peptide. We show that the addition of SnoopTag does not alter the secondary structure of the RALA domain which is essential for transfection and endosomal escape.^25^ Then we demonstrate that Snoop-RALA can be integrated into nanoparticles at up to 20% of the total RALA molecules while maintaining a nanoparticle size less than 150 nm and charge from 20-30 mV, both of which are critical parameters for the endocytosis and uptake of nanoparticles.^75^ Size and charge measurements also directly correlated with transfection efficiency with the transfection yields constant with up to 20% Snoop-RALA. The reduced transfection percentage and improved cell viability also indicates that SnoopTag residues are present on the outside of nanoparticles, altering the surface properties. This was further demonstrated by the efficient binding of the Catcher protein to the surface of the nanoparticles.

With the functionalized nanoparticle system in place, we selected zwitterionic SBMA microgels as the foundation for nanoparticle binding due to its reduced non-specific protein adsorption and its capacity to support MSC proliferation and differentiation.^55,56^ We add SpyTag and RGD peptides to the surface of these microgels, then demonstrate that the Catcher protein covalently tethers Snoop-RALA nanoparticles to the microgel surface. This system allowed for the sustained release of the nanoparticles over a two week span with less than 25% of the nanoparticles released. This release profile establishes the strength of covalent docking of nanoparticles and enables the future incorporation of enzyme cleavable peptides for fine control over the release kinetics in response to physiological signals for stimuli-responsive release. For example, MMP2 cleavable peptides could be integrated to release nanoparticles in response to inflammation, halting tissue degradation and guiding tissue repair in immune disorders such as osteoarthritis (OA).^76^

Using our RALA-Catcher-microgel system we demonstrate prolonged delivery of mRNAs for sustained gene expression in MSCs seeded within a jammed microgel scaffold. Analysis of nanoluc expression demonstrates that mRNA expression can be extended for over 21 days with minimal differences in the presence or absence of the Catcher protein linker. Similarly, RT-qPCR demonstrates sustained CRISPR activation of *PRG4*. Expression was significantly upregulated through day 7, whether nanoparticles were tethered or untethered, and remained over 6-fold greater than non-transfected controls at day 14, though this difference was no longer significant relative to non-targeting controls. These results indicate that nanoparticles are trapped within the packed microgel scaffold regardless of covalent tethering. Interestingly, untethered nanoparticles resulted in significantly higher *PRG4* activation on day 7 and also trended higher on day 14, which may indicate that covalent tethering restricts nanoparticles from accessing the cells, limiting the overall transfection despite the sustained release curve. Although these results were not reflected in our nanoluc expression data, we previously demonstrated that CRISPR activity is more reliant on higher transgene copy numbers compared to reporter mRNAs.^14^ These results demonstrate the importance of adding cell-cleavable linkers to allow for nanoparticle separation from the microgels and the sustained uptake of CRISPR cargos.

Despite these limitations, our biomaterial system sustained *PRG4* activation for over double the duration of monolayer transfections with much greater *PRG4* fold-changes. Although direct comparison across studies is difficult due to differences in normalization approaches, target gene selection, and cell type, our system demonstrates sustained gene activation over 6-fold relative to untransfected controls at day 14, comparable to previous CRISPRa-biomaterial systems^42^. Importantly, unlike these prior reports, our study includes non-targeting controls to distinguish gene-specific activation from transfection-associated effects, providing a more accurate assessment of CRISPRa activity.^42^ We demonstrate that this influence of transfection on target gene expression is especially important in 3D transfection as seen by the variable *PRG4* fold-changes with our non-targeting groups compared to the constant downregulation of *PRG4* in monolayer non-targeting groups. These results highlight a key consideration that cells have unique unintended responses to transfection in 3D zwitterionic microgels compared to 2D tissue culture plastic. In fact, Truong *et al*. previously demonstrated that gene transfer in 3D microporous annealed scaffolds relied on caveolae-mediated endocytosis rather than clathrin-mediated endocytosis that is more dominant in 2D culture.^77^ Transcriptomic analysis of RALA transfection in 3D microgel scaffolds should be performed to identify further unexpected transcriptomic effects, especially those linked with inflammatory responses that may hinder the downstream impact of CRISPR gene editing.

While this system represents a promising foundation for spatiotemporal regulation of CRISPR gene editing, there are many opportunities to build upon this framework due to its high tunability. We demonstrate that nanoparticles containing up to 20% Snoop-RALA nanoparticles retain their transfection capacity, leaving many available SnoopTag moieties after nanoparticle binding to microgel scaffolds. Since the SpyTag system is well established for adding cell-specific ligands or therapeutic proteins to nanoparticles, future work should investigate further decorating nanoparticles for cell-targeting, increased retention, or other specific applications.^50^ However, in doing so, it is important to carefully titrate the amounts of fusion proteins, since overloading protein content on the nanoparticle surface can inhibit endosomal escape and therefore transgene expression.^78^ Additionally, this system can be modified easily for dual CRISPR strategies such as dual activation and inhibition of specific genes for enhanced control over cell phenotype. Simply interchanging the system for SpyTag-functionalized RALA and SnoopTag decorated microgels will allow for localized binding of the respective nanoparticles using the same Catcher protein as the linker.

Overall, we present a highly versatile 3D microgel scaffold for spatiotemporal controlled CRISPR gene editing. We demonstrate that cell-friendly RALA nanoparticles can be modified with SnoopTag peptide to allow for efficient binding to zwitterionic microgels. Then, we show that jammed microgels allow for prolonged mRNA expression and sustained CRISPR activity. Future work must characterize the effects of this system over MSC function and lineage commitment and the influence of different CRISPR modalities on these processes. Furthermore, this work fabricates 3D biomaterial scaffolds by jamming microgels into a continuous scaffold. Compared to previous biomaterial-based CRISPR delivery strategies, microgels are a powerful platform for injectable biomaterials, capable of filling irregular defects, conforming to surrounding tissue, and supporting minimally invasive drug delivery and/or tissue regeneration. However, covalent linkages between microgels are required improve the clinical effectiveness of this approach. Multiple strategies have been developed to form microporous annealed particle (MAP) scaffolds. For example, Griffin *et al*. developed a system in which two peptides incorporated into hyaluronic acid microgels are enzymatically crosslinked by factor XIIIa, allowing for rapid annealing after injection.^79,80^ Similar systems could be incorporated alongside the SnoopTag/SpyTag system to improve scaffold stiffness and ensure precise localization after injection.

## 5. Conclusions

This work establishes a novel system for covalently tethering non-viral CRISPR nanoparticles to zwitterionic microgels for spatial and temporal control over gene editing. We integrate the SnoopTag peptide into RALA nanoparticles without impacting nanoparticle formation or transfection rates. Then, we demonstrate that these functionalized nanoparticles bind to zwitterionic microgels with high affinity via a SnoopCatcher-SpyCatcher linker protein. Finally, we harness this system to deliver mRNA to MSCs for sustained gene expression and CRISPR activity compared to monolayer transfections. This work lays the framework for a highly tunable platform for customizable gene editing strategies to engineer complex tissue gradients to benefit a plethora of therapeutic applications.

## 6. Acknowledgements

This material is based upon work supported by the National Science Foundation under Grant No. 2441592 to Tomas Gonzalez-Fernandez. Any opinions, findings, and conclusions or recommendations expressed in this material are those of the author(s) and do not necessarily reflect the views of the National Science Foundation. Additionally, Tomas Gonzalez-Fernandez would like to acknowledge the ON/ORS Kick-Starter Grant number 23-141 from the Orthoregeneration Network (ON) Foundation (Switzerland) and the Orthopaedic Research Society, the Career Development Award from the American Society of Gene and Cell Therapy, and the start-up funds provided by the Department of Bioengineering and the P.C. Rossin College of Engineering and Applied Science at Lehigh University. The content is solely the responsibility of the authors and does not necessarily represent the official views of the American Society of Gene and Cell Therapy. Josh Graham would like to acknowledge the support provided through the National Science Foundation Graduate Research Fellowship under Grant No 2234658.

