## Supplemental information for "Spatiotemporal Control of Genetic and Epigenetic Editing through Covalent Tethering of CRISPR Nanoparticles to Zwitterionic Microgels"

Joshua P. Graham[1], Erika E. Wheeler[2,3], J. Kent Leach[2,3], Tomas Gonzalez-Fernandez[1,4]
[1]Department of Bioengineering Lehigh University.
[2]Department of Biomedical Engineering, University of California Davis.
[3]Department of Orthopaedic Surgery, University of California Davis Health.
[4]Institute for Functional Materials and Devices, Lehigh University.

### Supplemental information

### Supplemental Tables

Table S1. *PRG4* gRNA sequence used in this study. 2

### Supplemental Figures

Figure S1. *PRG4* gRNA sequence optimization. 2
Figure S2. Snoop-RALA transfection profiles. 3

**Table S1.** gRNA sequences used in this study. Sequences in bold were used in the main text.

| Gene Symbol | Label | Sequence |
|---|---|---|
| *PRG4* | **A** | **GGGCCCAGACGACTAGACTT** |
| | B | GTGAACAAATTGTCCTGAGA |
| | C | GTATGCCACAAGCTAACACT |
| *AAVS1* | **A** | **GGGGCCACTAGGGACAGGAT** |

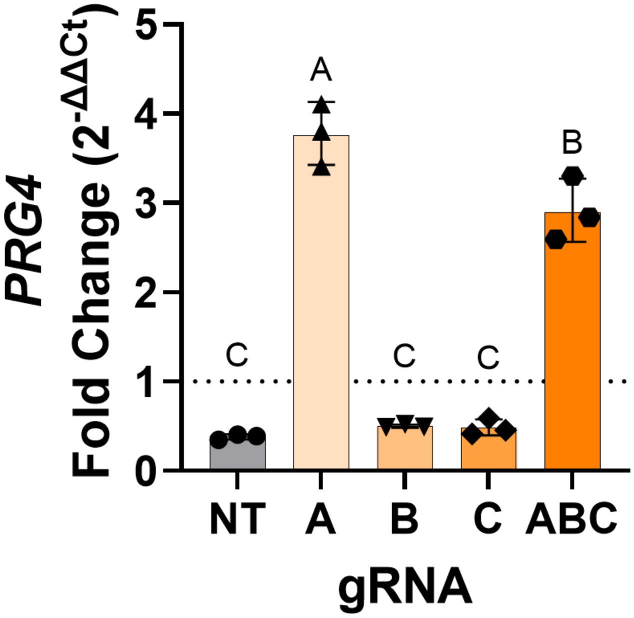


**Figure S1.** gRNA screening and optimizations for *PRG4* activation. Three gRNAs listed in Table S1 or non-targeting gRNA (NT) were delivered to MSCs in monolayer and analyzed via RT-qPCR relative to non-transfected controls. Data plotted as the geometric mean ± the geometric standard deviation (n = 3). Samples that are not statistically significantly different ($p < 0.05$) as determined by a one-way ANOVA are denoted by the same letter.

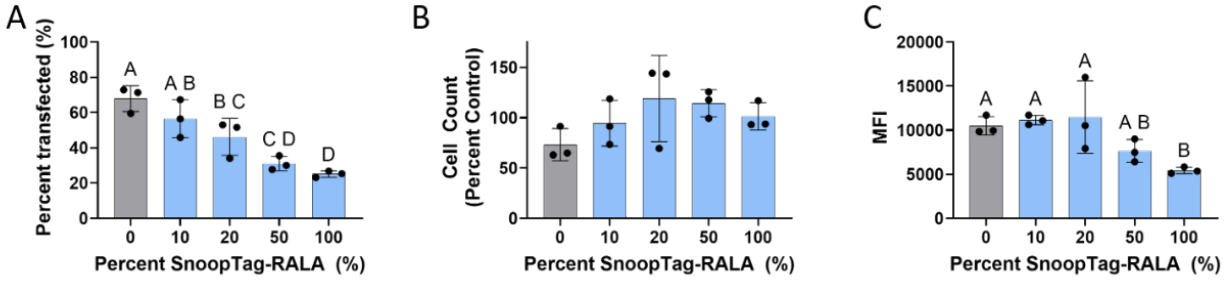


**Figure S2.** Snoop-RALA transfection profiles. SnoopTag functionalized RALA peptides were integrated into RALA nanoparticles at different molar percentages then used to transfect MSCs with GFP mRNA. **A)** The percentage of transfected cells, **B)** the cell viability represented by the cell counts as a percentage of the control. And **C)** the mean fluorescent intensity of transfected cell populations. Data plotted as mean ± standard deviation (n = 3). Samples that are not statistically significantly different ($p < 0.05$) as determined by a one-way ANOVA are denoted by the same letter.
